# Protein Age Bias in Target Degradation by PROTACs

**DOI:** 10.64898/2026.05.01.722262

**Authors:** Bingbing X. Li, Xiangshu Xiao

**Affiliations:** Program in Chemical Biology, Department of Chemical Physiology and Biochemistry, Knight Cancer Institute, 3181 SW Sam Jackson Park Rd, Portland, Oregon, USA

## Abstract

Targeted protein degradation (TPD) by PROteolysis TArgeting Chimeras (PROTACs) has emerged as a powerful chemical biology and therapeutic modality, yet many degraders exhibit incomplete target clearance and characteristic rebound kinetics despite continuous exposure. The mechanistic basis for this behavior remains poorly understood. Here we uncover protein age as a previously unrecognized determinant of PROTAC efficacy. Using _CG_□SLENP, a chemical genetics strategy that selectively labels newly synthesized and pre □existing proteins within the same living cell, we directly resolve PROTAC□induced degradation of distinct intracellular protein populations. Applying this approach to the bromodomain protein BRD4, we show that two mechanistically and structurally distinct PROTACs, dBET6 and MZ□1, preferentially degrade pre □existing BRD4, while newly synthesized BRD4 is degraded substantially more slowly and incompletely. This age□dependent degradation bias is observed in live□cell imaging, across compound concentrations and time scales, and for both reporter and endogenous BRD4. These findings reveal that PROTAC□mediated degradation is governed not only by target engagement and ternary complex formation, but also by the dynamic balance between protein synthesis and degradation. By identifying temporal proteostasis as a critical parameter in TPD, this work provides a mechanistic framework for incomplete degradation and rebound kinetics and establishes protein maturation state as an important consideration for degrader design and evaluation.

## Introduction

PROteolysis TArgeting Chimeras (PROTACs) are heterobifunctional molecules composed of a target-binding ligand, a E3 ligase ligand and a linker to connect them.^1^ PROTACs represent a promising therapeutic strategy for numerous diseases to achieve targeted protein degradation (TPD). Unlike traditional inhibitors, PROTACs offer selective elimination of target proteins rather than merely inhibiting their activities.^1–3^ Thus, a PROTAC can be designed to remove a target protein even when the ligand does not target the active site of the target protein. During the past decade, this approach has garnered significant attention in the scientific community for its potential in targeting traditionally undruggable proteins, overcoming drug resistance, and achieving prolonged pharmacodynamic effects with a catalytic mechanism of action.^4–10^ While the design of PROTACs remains largely empirical,^11–13^ relying heavily on trial-and-error optimization of linker length, E3 ligase recruitment, and ternary complex formation, the intensive investigations of PROTACs have led to the advance of over 40 PROTACs into the clinical arena for different maladies.^14, 15^

PROTACs eliminate the entire target protein through formation of ternary complexes containing target protein, E3 ligase and PROTACs themselves. This is followed by polyubiquitination and degradation by the ubiquitin proteasome system (UPS).^16^ It has been assumed that the mechanism of recovery depends on new protein synthesis. However, it has not been established if and how PROTACs degrade newly synthesized proteins. Literature reports showed that many PROTACs exhibit atypical degradation kinetics in which the target protein level initially declines but subsequently rebounds despite the continued presence of the degrader. This phenomenon was first observed with the bromodomain-containing 4 (BRD4)-targeting PROTAC dBET1, which induced robust BRD4 degradation during the first 16 hours, followed by substantial recovery by 24 hours under constant compound exposure.^17^ Similar rebound kinetics have been reported for other BRD4 degraders,^18^ as well as for androgen receptor (AR) degraders such as ARV-110 and the structurally distinct ARD-1676.^19, 20^ The time course experiment with a fixed concentration of BRD9 degrader DBr-1 also showed rapid degradation after 30□min exposure and a rebound after 24□h.^21^ The mechanisms underlying this unusual degradation kinetics remain elusive.

Compared to existing proteins, newly synthesized proteins can assume different posttranslational modifications and also participate in different protein complexes. For example, the newly synthesized crystallins in the eye lens are properly folded and highly soluble up to >450 mg/mL.^22^ Once synthesized, crystallins are retained in the lens for the entire decades of human lifespan. As we age, the crystallins get oxidized and become aggregates to form cataract affecting vision.^22^ In another example, the plasma protein apolipoprotein A-1 (ApoA-1) has been shown to undergo asparagine/glutamine deamination over protein aging.^23^ This protein deamination is facilitated in patients with type 1 diabetes, which can be normalized to a certain extent with insulin treatment.^23^ Besides these individually studied proteins, measurements at the whole proteome scale revealed that newly synthesized proteins display faster degradation kinetics than old proteins, likely reflecting different protein complexes are invovled.^24^ Together, these studies have clearly demonstrated that newly synthesized proteins and pre-existing proteins are not the same. However, the challenge to differentiate newly synthesized proteins from existing proteins in the same cells is daunting. To address this challenge, we recently reported _CG_-SLENP, a <u>c</u>hemical <u>g</u>enetics-based approach for <u>s</u>elective labeling of <u>e</u>xisting and <u>n</u>ewly synthesized <u>p</u>roteins,^25,26^ that enables selective labeling and visualizing newly synthesized and pre-existing proteins in live cells. With _CG_-SLENP, we showed that small molecule lamin inhibitor **LBL1** differentially modulates the localization and assembly of newly synthesized and pre-existing nuclear lamin A in live cells.^25^

In this article, we employed _CG_-SLENP to investigate the degradation of profiles of newly synthesized and existing BRD4 in live cells with different BRD4-targeting PROTACs. These investigations show that newly synthesized BRD4 is less susceptible to degradation induced by different PROTACs. This differential susceptibility suggests that incomplete clearance of newly synthesized protein may contribute to the limited efficacy and characteristic target protein rebound kinetics observed for multiple PROTACs.

## Results and Discussions

### _CG_-SLENP to label existing and newly synthesized BRD4

In order to investigate the degradation effects of PROTACs on existing BRD4 and newly synthesized BRD4, we developed _CG_-SLENP to label existing and newly synthesized BRD4. To this end, a lentiviral construct encoding a Flag tagged HaloTag fused in-frame with BRD4 was generated and sequence-verified (see Figure S1 for sequence). HaloTag is a self-labeling protein tag that covalently reacts with 6-chlorohexanyl derivatives with minimal background labeling in mammalian cells.^25, 27^ MDA-MB-468 cells were transduced with this lentivirus to achieve stable expression of HaloTag-BRD4. As shown in Figure 1B, the fusion protein was expressed at a level comparable to that of endogenous BRD4. To ensure the expressed HaloTag-BRD4 functioned properly, we investigated the localization of exogenously expressed HaloTag-BRD4 using immunofluorescence. Similar to endogenous BRD4, the exogenously expressed HaloTag-BRD4 was also exclusively localized in the nucleus (Figure 1C). The staining pattern of anti- Flag (M2) was indistinguishable from that of BRD4. To determine if the exogenously expressed HaloTag-BRD4 could be labeled by a Halo ligand, we treat the cells with Halo-JF549 (Figure 1A). As shown in Figure 1D, HaloTag-BRD4 was efficiently labeled with Halo-JF549. The pattern of Halo-labeling was the same as that of M2 staining. Furthermore, neither Halo labeling nor M2 staining was observed in parental MDA-MB-468 cells (Figure 1C-D), supporting the specificity of Halo labeling. Together, these results demonstrate that HaloTag-BRD4 is functional and can be selectively labeled using Halo ligands.

**Figure 1.**
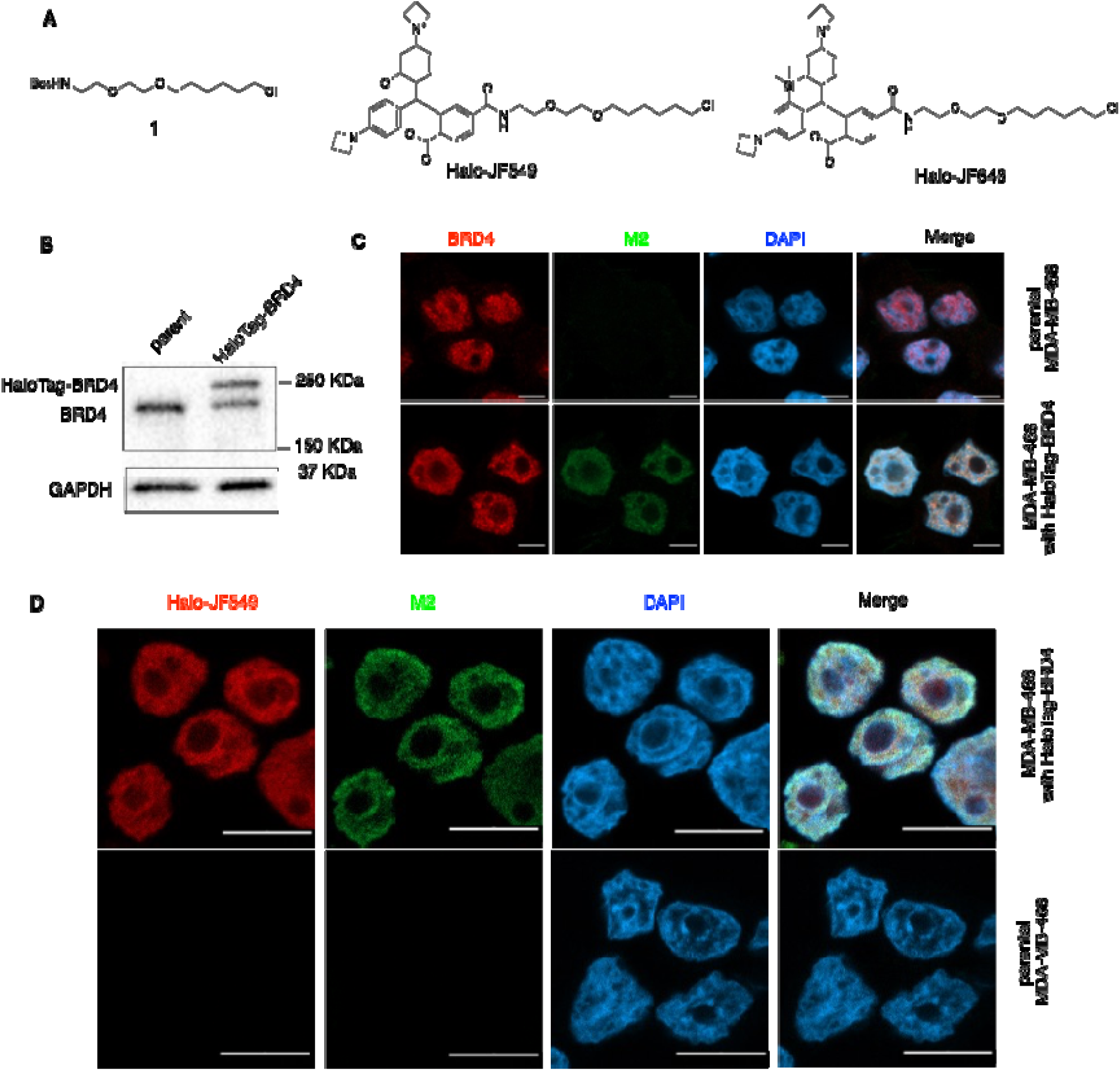
Expression of HaloTag-BRD4 in MDA-MB-468 cells. (A) Chemical structures of different Halo ligands used in this article. (B) Western blot analysis of HaloTag-BRD4. The lysates from parental MDA-MB-468 cells and cells transduced with HaloTag-BRD4 were prepared for western blotting analysis using anti-BRD4 and anti-GAPDH. (C) HaloTag-BRD4 was localized in the nucleus. MDA-MB-468 cells transduced with HaloTag-BRD4 and parental MDA-MB-468 cells were fixed, permeabilized and stained with indicated antibodies for confocal analysis. The nuclei were countered stained with DAPI. (D) Halo labeling of HaloTag-BRD4. The indicated cells were treated with Halo-JF549. Then the cells were fixed, permeabilized and stained with indicated antibodies for confocal analysis. The scale bars are 10 μm.

With the HaloTag-BRD4 system established, we investigated if the existing and newly synthesized HaloTag-BRD4 could be selectively labeled using _CG_-SLENP. To label existing HaloTag-BRD4, MDA-MB-468 cells with HaloTag-BRD4 were treated with a protein synthesis inhibitor cycloheximide (CHX) along with Halo-JF549 (Figure 2A). As shown in Figure 2B, existing HaloTag-BRD4 was efficiently labeled with Halo-JF549, similar to the results shown in Figure 1D. To label newly synthesized HaloTag-BRD4, we resorted to a silent Halo ligand **1** (Figure 1A).^25^ The cells were first treated with ligand **1** to label existing HaloTag-BRD4, which became invisible under a confocal microscope (Figure 2C). Upon removal of excess ligand **1**, the cells were allowed to continue the synthesis of new proteins. At different time points of recovery of protein synthesis, the cells were then treated with Halo-JF549 to label any newly synthesized HaloTag-BRD4. Within the first 15 min after removal of ligand **1**, no Halo-JF549 was detected (Figure 2D, 0, 15 min), demonstrating complete labeling of existing HaloTag-BRD4 by **1** and newly synthesized HaloTag-BRD4 was minimal. Around 30 min post recovery, visible but weak Halo-JF549 signal was observed. At 60 or 120 min post recovery, prominent newly synthesized HaloTag-BRD4 was clearly observed (Figure 2D). To demonstrate that the observed Halo signal was indeed due to newly synthesized HaloTag-BRD4, we treated the cells with CHX after labeling with ligand **1** (Figure S2A). At the different time points (0-120 min) post labeling with ligand **1**, addition of Halo-JF549 did not produce any detectable Halo signals, supporting that the new Halo signal observed in Figure 2D was from newly synthesized HaloTag-BRD4, rather than existing HaloTag-BRD4 that re-exposed its binding site during the recovery phase. Together, these result demonstrate that _CG_-SLENP can be used to selectively label existing HaloTag-BRD4 and newly synthesized HaloTag-BRD4.

**Figure 2.**
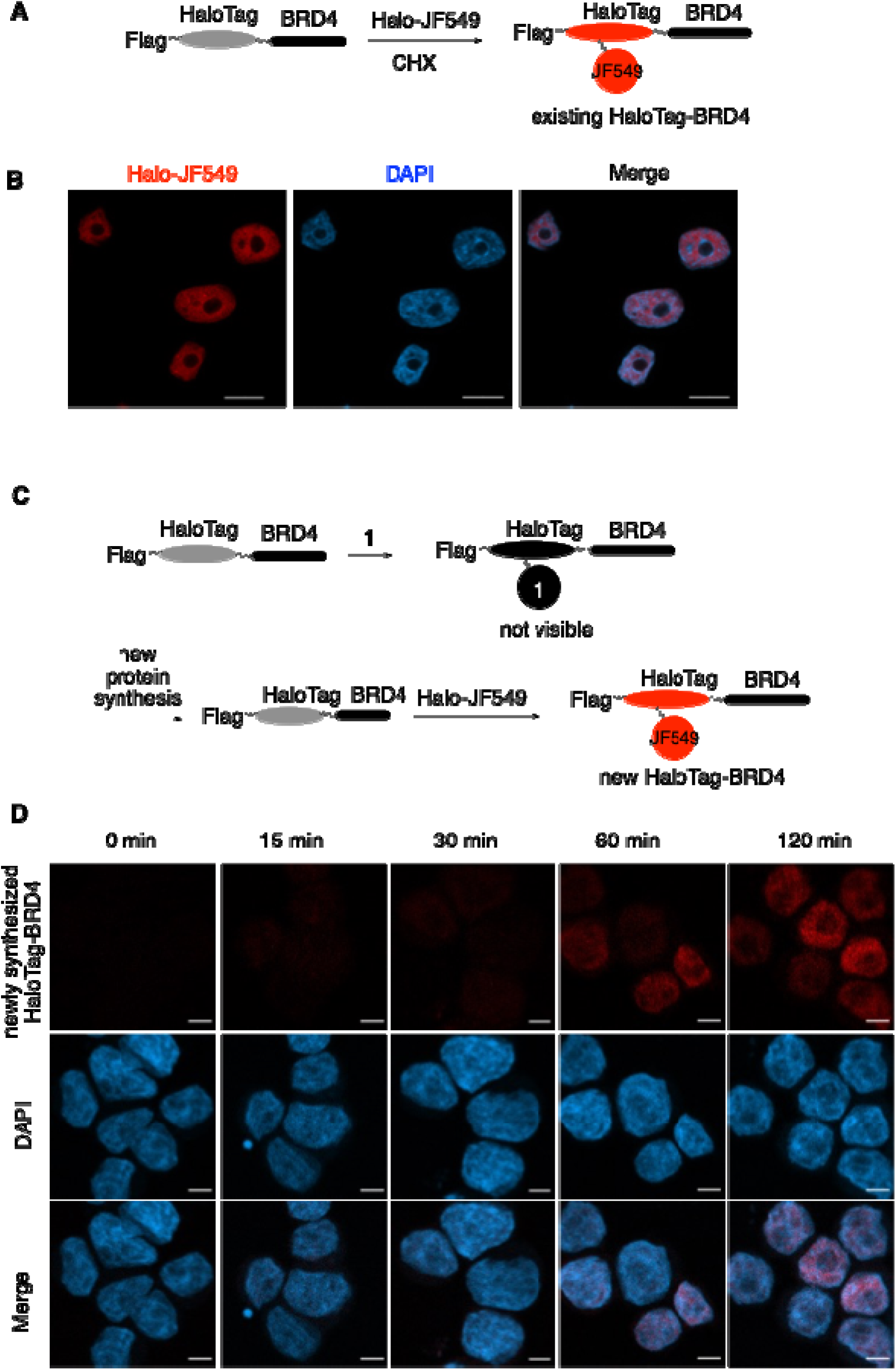
_CG_-SLENP for selectively labeling existing and newly synthesized HaloTag-BRD4. (A) Experimental design to label existing HaloTag-BRD4. (B) Representative images of existing HaloTag-BRD4 labeled by Halo-JF549. MDA-MB-468 cells with HaloTag-BRD4 were treated with Halo-JF549 in the presence of CHX for 30 min. The nuclei were counter stained with DAPI. (C) Experimental design to label newly synthesized HaloTag-BRD4. (D) Representative images of newly synthesized HaloTag-BRD4 labeled by Halo-JF549. MDA-MB-468 cells with HaloTag-BRD4 were treated with Halo ligand **1** for 30 min. Then the cells were allowed to synthesize new protein in the absence of CHX for different periods of time, when Halo-JF549 was added for another 10 min. The nuclei were counter stained with DAPI. Scale bars are 5 μm.

### _CG_-SLENP to dually label existing and newly synthesized HaloTag-BRD4 in the same cells

Having verified that _CG_-SLENP could be harnessed to selectively label existing and newly synthesized HaloTag-BRD4, we asked if _CG_-SLENP could be employed to dually label both existing and newly synthesized HaloTag-BRD4 in the same cell, which would be highly advantageous to investigate the potential degradation differences by PROTACs. To this end, the cells were first labeled with Halo-JF646 (Figure 1A) to label existing HaloTag-BRD4. Then the excess Halo ligand was removed through extensive washings, after which the cells were allowed to resume new protein synthesis. At different time points post resumption of new protein synthesis, the cells were further treated with Halo-JF549 to label newly synthesized HaloTag-BRD4 (Figure 3A). The dual labeling results are shown in Figure 3B. Consistent with the results shown in Figure 2B, the existing HaloTag-BRD4 was efficiently labeled with Halo-JF646. No newly synthesized HaloTag-BRD4 (red signal) was detected immediately after Halo-JF646 labeling (0 min). By 30 min, faint Halo-JF549 signals representing newly synthesized HaloTag-BRD4 appeared. By 1 hour and beyond, robust labeling of newly synthesized HaloTag-BRD4 was observed. In each cell, the distribution pattern of the newly synthesized HaloTag-BRD4 was indistinguishable from that of the existing one, suggesting that the newly synthesized HaloTag-BRD4 is rapidly translocated to the nucleus without being retained in the cytosol. These findings were consistent with the results shown in Figure 2.

**Figure 3.**
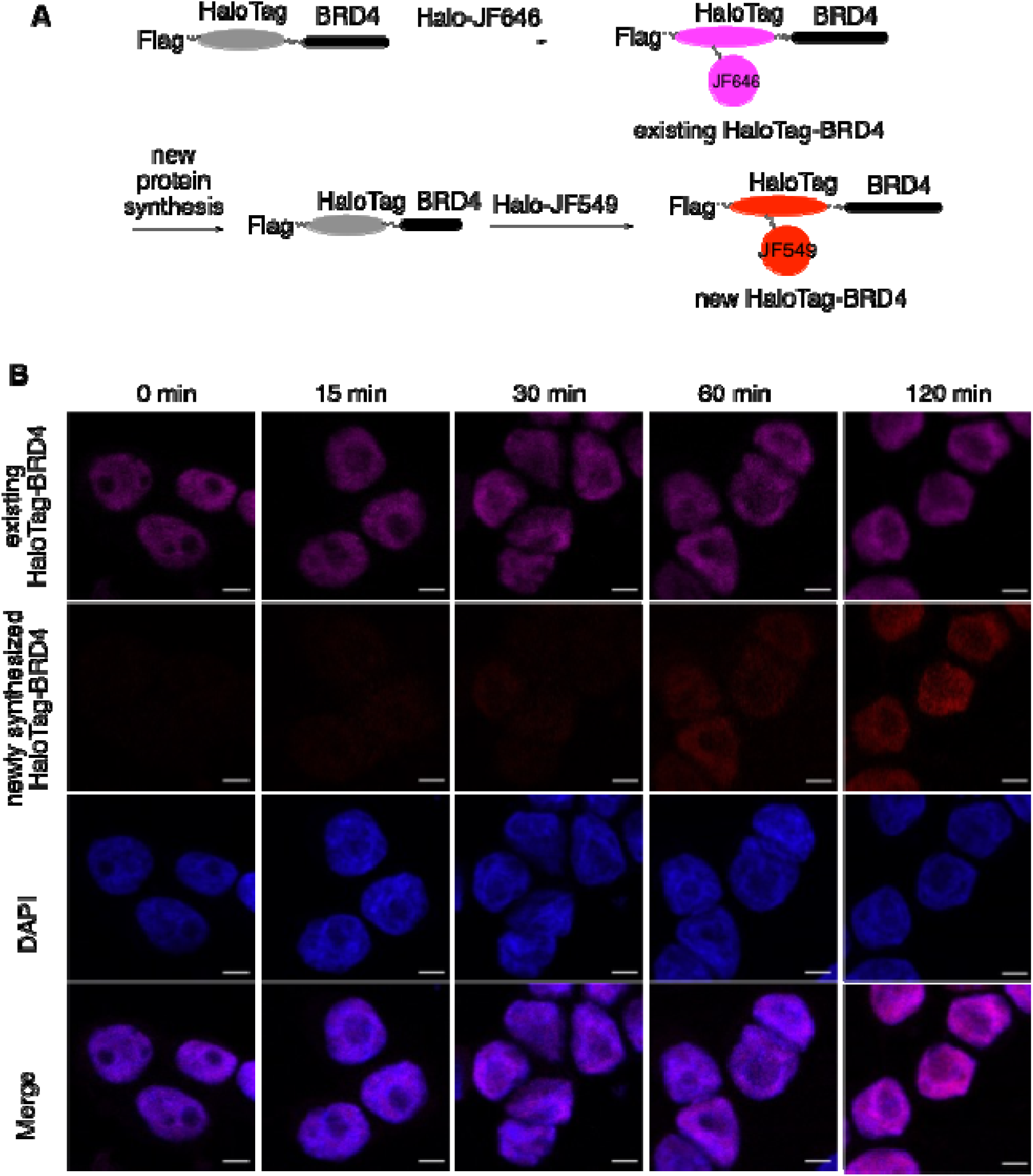
_CG_-SLENP for dual labeling of existing and newly synthesized HaloTag-BRD4. (A) A schematic diagram to illustrate the dual-labeling _CG_-SLENP strategy to label existing and newly synthesized HaloTag-BRD4 in the same cells. (B) Representative confocal micrographs of existing and newly synthesized HaloTag-BRD4. MDA-MB-468 cells expressing HaloTag-BRD4 were treated with Halo-JF646 for 1 h, when the ligand was removed. The cells were incubated in drug-free media for the indicated time period, when the cells were treated with Halo-JF549 for 15 min. The cells were then fixed and permeabilized. The cell nucleus was counterstained with DAPI. Scale bars are 5 μm.

To confirm that Halo-JF549 only labeled newly synthesized HaloTag-BRD4, the cells were treated with CHX to inhibit new protein synthesis after labeling with Halo-JF646 (Figure S3A). As shown in Figure S3B, the existing HaloTag-BRD4 was efficiently labeled by Halo-JF6464. However, no Halo-JF549 signal was detected at any time points post Halo-JF646 labeling (0-120 min). These results confirmed that the red signal of Figure 3B indeed represented newly synthesized HaloTag-BRD4. Therefore, _CG_-SLENP provides an effective way to selectively label existing and newly synthesized HaloTag-BRD4 in the same cells.

### dBET6 preferentially degrades existing BRD4 over newly synthesized BRD4

Having established that _CG_-SLENP could be utilized for dually labeling of existing and newly synthesized HaloTag-BRD4 in the same cells, we began to ask if a BRD4-targeting PROTAC had differential effect on the target degradation. To investigate this notion, we selected dBET6 as a model degrader. dBET6 is a potent BRD4-targeting PROTAC constructed by conjugating BRD4 inhibitor JQ1 with a CRBN ligand.^28^ In MDA-MB-468 cells, we found that dBET6 efficiently degraded BRD4 after 24 h treatment (Figure S4). However, upon prolonged treatment for 48 or 72 h, we found that the degradation efficiency was greatly diminished. This rebound kinetics is similar to other reported PROTACs in different cellular contexts.^17–21^

We used _CG_-SLENP approach to dually label existing and newly synthesized HaloTag-BRD4 as shown in Figure 4A. Specifically, the existing protein was labeled with Halo-JF549 while the newly synthesized protein was labeled with Halo-JF646. Once both pools were labeled, we treated the cells with dBET6 and monitored the degradation of existing (Halo-JF549) and newly synthesized (Halo-JF646) pools for 3 h using live cell imaging. The fluorescence intensities of the existing and newly synthesized HaloTag-BRD4 were then extracted and quantified on a per cell basis. In the absence of dBET6, a small reduction of both existing and newly synthesized HaloTag-BRD4 was observed (Figure 4B). Interestingly, the basal degradation of existing HaloTag-BRD4 was slightly higher than that of the newly synthesized one (Figure 4D, *P* < 0.01). When the cells were treated with dBET6 (10 nM), the existing HaloTag-BRD4 was rapidly degraded and only 35% of HaloTag-BRD4 remained at 3 h post dBET6 treatment (Figure 4C-D). On the other hand, the degradation of newly synthesized HaloTag-BRD4 was much more sluggish and as much as 72% of newly synthesized HaloTag-BRD4 remained. While the basal degradation rate of newly synthesized HaloTag-BRD4 was slower than that of the existing one, this time-dependent divergence in degradation between new and existing HaloTag-BRD4 induced by dBET6 was still significant (Figure 4D, *P* <0.01). To investigate dose-dependence of this differential degradation by dBET6, we treated the cells with a series of different concentrations of dBET6 for live cell imaging after the existing and newly synthesized HaloTag-BRD4 were dually labeled. As shown in Figure 4E, the ratio of newly synthesized to existing HaloTag-BRD4 was increased over time, consistent with the conclusion that existing HaloTag-BRD4 was preferentially degraded. The treatment exhibited a dose-dependent response (10 nM > 5 nM > 1 nM) that became progressively more pronounced over the course of the experiment of 180 min (Figure 4E).

**Figure 4.**
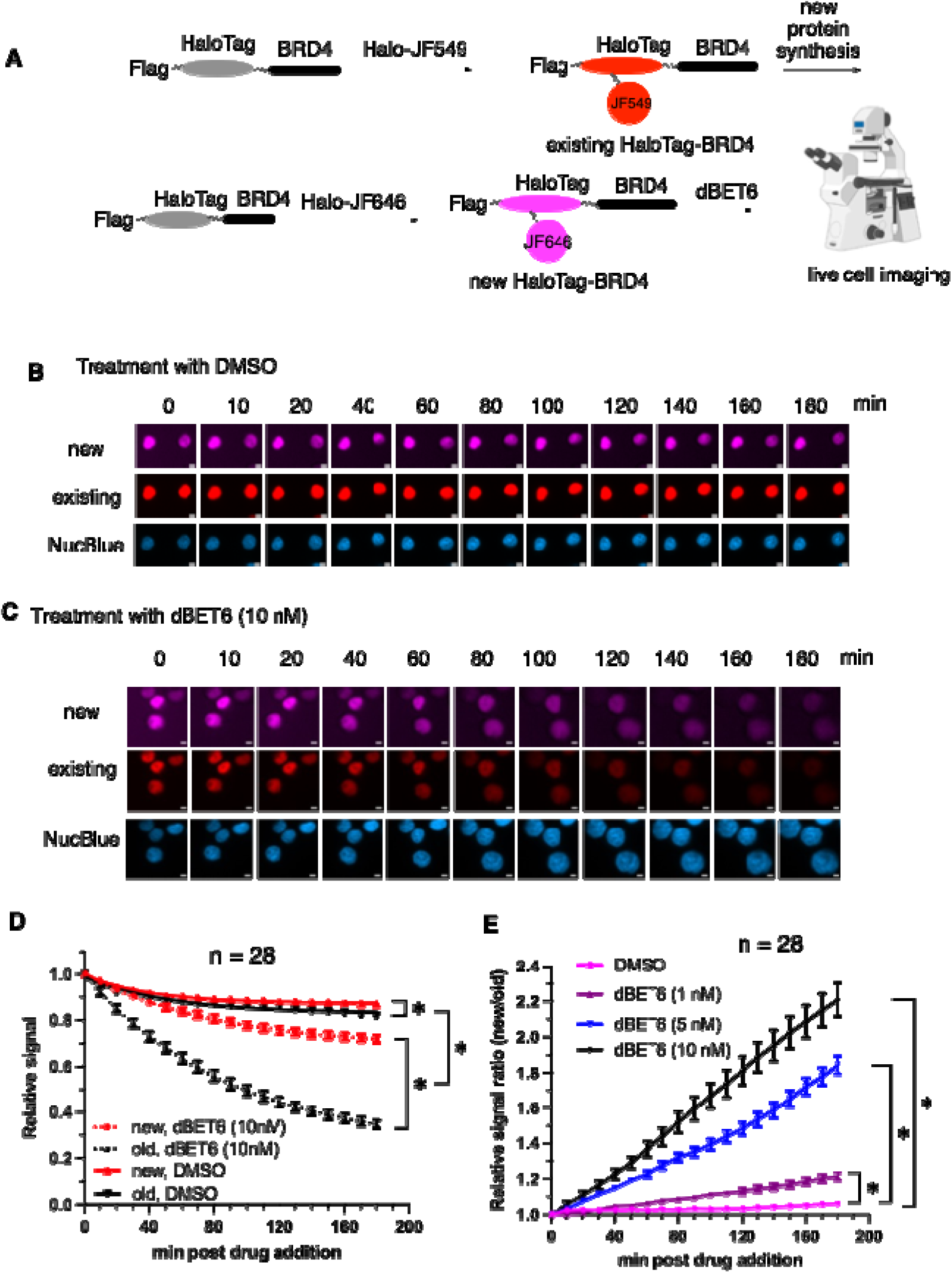
dBET6 preferentially degrades existing HaloTag-BRD4 over newly synthesized HaloTag-BRD4. (A) A schematic diagram to illustrate dual-labeling by _CG_-SLENP to investigate the degradation effects of dBET6 on existing and newly synthesized HaloTag-BRD4 in live cells. (B, C) Representative live cell micrographs of cells treated with DMSO (B) or dBET6 (10 nM, C). Existing HaloTag-BRD4 was labeled with Halo-JF549 while the newly synthesized HaloTag-BRD4 was labeled with Halo-JF646 (far-red). Upon labeling, the cells were treated with DMSO or different concentrations of dBET6 for live-cell imaging. The images were acquired every 10 minutes for 3 hours in the red, far-red, and NucBlue channels. Scale bars are 5 μm. (D) Time-dependent degradation of HaloTag-BRD4 by dBET6. The existing and newly synthesized HaloTag-BRD4 signals from images in (B) and (C) were quantified. Data are presented as mean ± SEM (n = 28). \**P*<0.01. (E) dBET6 dose-dependently induced preferential degradation of existing HaloTag-BRD4. The data in (D) were replotted by dividing newly synthesized HaloTag-BRD4 to the existing HaloTag-BRD4. These divided values were normalized to 1.0 at time zero. Data are presented as mean ± SEM (n = 28). \**P*<0.01.

The results presented in Figure 4 are the first time to show that an existing protein is preferentially degraded over newly synthesized protein by a PROTAC in the same cell. We were curious if this effect could be extended to other PROTACs. Thus we investigated MZ-1, a BRD4-targeting PROTAC by engaging VHL, another commonly employed E3 ligase for PROTAC development.^29^ The existing and newly synthesized HaloTag-BRD4 was similarly labeled as shown in Figure S5A. Then the cells were treated with MZ-1 at different concentrations for live cell imaging for 3 h. Similar to dBET6, MZ-1 also preferentially degraded existing HaloTag-BRD4 (Figure S5B-S5E). This preference was both dose-dependent and becoming more pronounced at later time points (Figure S5D-S5E). Together with the dBET6 data, these findings indicate that time-dependent and dose-dependent divergence of degradation of new and existing BRD4 may be a generalizable feature of BRD4-directed PROTACs.

### dBET6 preferentially degrades existing endogenous BRD4 over newly synthesized endogenous BRD4

Having demonstrated selective degradation of existing HaloTag-BRD4 by PROTACs in our HaloTag-reporter system using _CG_-SLENP, we next assessed whether this effect was translatable to the endogenous BRD4 protein. We took advantage of the fact that dBET6 is able to degrade existing BRD4, and BRD4 protein is continuously being synthesized in the cells. To investigate the effect of dBET6 on existing BRD4, MDA-MB-468 cells were treated with CHX along with dBET6 (Figure 5A, top). Then the remaining BRD4 level was assessed by immunofluorescence labeling. As shown in Figure 5B, all concentrations of dBET6 led to efficient BRD4 degradation. Quantification of residual BRD4 fluorescence intensity showed a robust dose-dependent degradation of existing BRD4 by dBET6 (Figure 5C).

**Figure 5.**
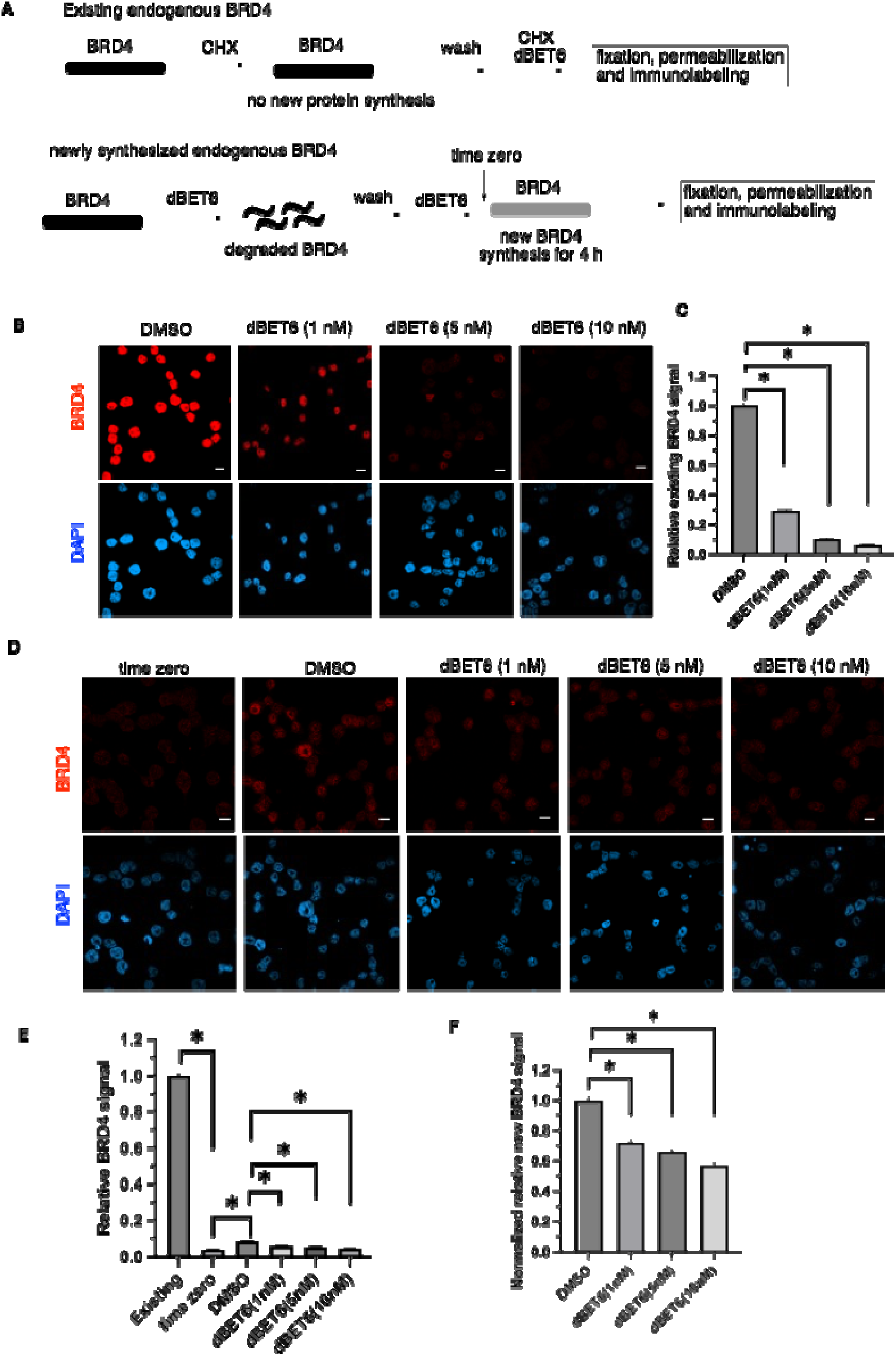
dBET6 preferentially degrades existing endogenous BRD4 over newly synthesized endogenous BRD4. (A) Schematic experimental design to examine dBET6-induced degradation of existing and newly synthesized endogenous BRD4 in MDA-MB-468 cells. (B) Representative fixed-cell confocal micrographs of degradation of existing BRD4 by dBET6. The cells were first treated with CHX for 4 h. Then the cells were treated with different concentrations of dBET6 along with CHX for another 4 h. The cells were fixed, permeabilized and immunostained with anti-BRD4. The nuclei were counter stained with DAPI. (C) The BRD4 fluorescent intensities in (B) were quantified. For each condition, 100-150 cells were counted. (D) Representative fixed-cell confocal micrographs of degradation of newly synthesized BRD4 by dBET6. The cells were first treated with dBET6 (100 nM) for 4 h to degrade existing BRD4. The residual dBET6 was removed by extensive washings. Then the cells were allowed to synthesize new proteins in the presence of different concentrations of dBET6 for 4 h, when the cells were fixed, permeabilized and immunostained with anti-BRD4. The nuclei were counter stained with DAPI. (E) The BRD4 fluorescent intensities in (D) were quantified. The existing BRD4 signal was from the DMSO-treated sample in (B). For each condition, 100-150 cells were counted. (F) Normalization of newly synthesized BRD4 signal in (E). The DMSO-treated sample in (E) was normalized to 1.0. All data are presented as mean ± SEM. \**P* < 0.01 by student *t*-test. Scale bars are 10 μm.

To directly assess the effect of dBET6 on newly synthesized BRD4 in MDA-MB-468 cells, the cells were first treated with relative high concentration of dBET6 (100 nM) for 4 hours to deplete existing BRD4. After removing the residual dBET6, the cells were allowed to synthesize new BRD4 for 4 h in the presence or absence of low concentrations of dBET6 (Figure 5A, bottom). The remaining BRD4 level was quantified by immunofluorescence. The initial 100 nM dBET6 treatment led to 96% reduction of the original BRD4 (time zero in Figure 5D and 5E). When the cells were allowed to recover for 4 h in the absence of dBET6, new BRD4 was synthesized and quantitative immunofluorescence analysis showed a 2-fold increase of BRD4 (Figure 5D–5E). When low concentrations of dBET6 were added, it did produce a dose-dependent suppression of BRD4 re-accumulation (Figure 5D–5E), suggesting newly synthesized BRD4 was also degraded by dBET6. However, the magnitude of degradation by dBET6 was much smaller compared to that of the existing BRD4. Specifically, dBET6 only degraded newly synthesized BRD4 to 72% at 1 nM, 66% at 5 nM and 57% BRD4 at 10 nM (Figure 5F). On the other hand, the existing BRD4 was degraded to 29% at 1 nM, 10% at 5 nM and 6% at 10 nM (Figure 5C). Together, these results demonstrate that dBET6 not only preferentially degraded existing HaloTag-BRD4, but also the endogenous existing BRD4.

## Discussions and Conclusions

Targeted protein degradation by PROTACs has emerged as a catalytic pharmacological strategy capable of eliminating disease□associated proteins that are refractory to conventional inhibition. Among the many advanced PROTACs, an estrogen receptor (ER)-targeting PROTAC vepdegestrant has demonstrated clinical superiority in *ESR1* mutant breast cancer patients in a Phase III trial,^30^ supporting the promise of this novel therapeutic modality. A central assumption underlying PROTAC function is that target recovery following degrader treatment occurs primarily through *de novo* protein synthesis after degradation of the pre□existing protein pool. However, it has been unknown how the newly synthesized proteins will be impacted by PROTACs. In this regard, many PROTAC degraders exhibit atypical pharmacodynamic behavior in which target proteins initially undergo robust depletion but subsequently re□accumulate despite continuous compound exposure.^17–21^ Although this rebound phenomenon has been widely reported across multiple degrader classes, including BRD4, AR, and BRD9□targeting PROTACs, its mechanistic basis has remained incompletely understood. In some cases, the chemical instability of PROTACs in-question (e.g. dBET1) can be a contributing factor.^31^

In the present study, we used the _CG_□SLENP chemical genetics platform to temporally resolve PROTAC□ induced degradation at the level of distinct intracellular protein populations in the same cells. By selectively labeling existing and newly synthesized BRD4 within the same living cell, we demonstrate that two mechanistically distinct BRD4□targeting PROTACs, dBET6 and MZ□1, preferentially degrade pre□ existing BRD4 while newly synthesized BRD4 is degraded much more slowly and incompletely. This divergence was consistently observed across time courses, compound concentrations, HaloTag□ BRD4 reporter and endogenous BRD4 systems, indicating that preferential degradation of the existing protein pool may represent an intrinsic property of PROTAC□mediated degradation rather than an artifact of exogenous expression. These findings suggest that degradability is not just a fixed biochemical property of a protein but also a dynamic function of its maturation state.

The exact mechanistic basis for this degradation divergence is currently unknown. However, a few possibilities exist. Newly synthesized BRD4 likely differs from pre□ existing BRD4 in several important respects, including post□translational modification status, chromatin engagement, or incorporation into higher□order transcriptional complexes. Such differences may influence the accessibility of lysine residues for ubiquitination or alter the conformational compatibility required for productive ternary complex formation. In this model, mature chromatin□associated BRD4 may represent a preferred substrate for PROTAC□induced ubiquitination, whereas nascent BRD4 may transiently occupy folding intermediates or protective protein assemblies that reduce encounter rates with the PROTAC-E3 ligase complex. Importantly, the preferential degradation of existing BRD4 observed here was recapitulated at the endogenous protein level. Acute depletion of BRD4 using high□dose dBET6 followed by re□ expression assays revealed that low nanomolar concentrations of dBET6 continued to suppress accumulation of newly synthesized BRD4, but to a substantially lesser extent than their degradation of the pre□ existing BRD4 pool. These results indicate that PROTAC efficacy may be constrained not only by target engagement and ternary complex formation but also by the continual production of nascent protein that may transiently escape degradation. Together, these findings introduce temporal proteostasis as a previously unrecognized determinant of targeted protein degradation.

In contrast to the traditional occupancy□driven pharmacology, where drug efficacy depends primarily on ligand binding affinity and residence time, PROTAC□mediated degradation appears to be influenced by the dynamic balance between protein degradation and protein synthesis. Under conditions of rapid target resynthesis, such as those observed for transcriptional regulators with short half□lives, newly synthesized protein may repopulate the cellular pool faster than it can be efficiently ubiquitinated, resulting in incomplete depletion or rebound kinetics despite continuous degrader exposure. In this regard, it is of interest to note that many PROTACs do not achieve maximal degradation (*D*_max_) of 100%,^32^ where the residual target proteins may be from newly synthesized proteins. Indeed, some partial degraders can induce complete degradation in the presence of CHX.^33^ Previous studies have shown that PTM patterns can vary as proteins age, with differences emerging over time scales of days to years.^22, 23^ We also recently reported that newly synthesized lamin A present different assembly state at the translating ribosomes within the time scale of minutes to hours.^25^ Future studies should be directed towards the understanding of the underlying mechanisms of the temporal target protein degradation preference by PROTACs. It is also of interest to determine if PROTACs using other newly developed E3 ligases^34^ or ligase-independent mechanisms^35, 36^ also exhibit similar behavior.

The protein maturation□dependent degradation has important implications for degrader design and clinical deployment. First, it suggests that linker optimization or E3 ligase selection alone may be insufficient to overcome incomplete degradation if nascent protein populations remain refractory to degradation. Second, it raises the possibility that rational combinations of degraders with agents that modulate protein synthesis, folding, or complex assembly may enhance target clearance by synchronizing the degradable state of the target protein. Finally, these findings highlight the importance of considering protein age or maturation state when evaluating degrader efficacy using bulk biochemical or endpoint assays, which may obscure differential degradation across intracellular protein subpopulations. More broadly, our results suggest that targeted protein degradation is governed not only by molecular recognition events but also by the temporal evolution of the target proteome. By enabling selective interrogation of newly synthesized and pre□existing protein pools in live cells, _CG_□SLENP provides a powerful platform for dissecting these dynamics with molecular precision. Application of this approach to additional degrader targets may reveal whether preferential degradation of mature protein represents a generalizable feature of PROTAC pharmacology.

In summary, we demonstrate that BRD4□directed PROTACs preferentially degrade pre□existing BRD4 over newly synthesized BRD4 in live cells. This differential susceptibility provides a mechanistic framework for understanding variable degradation efficiency and rebound kinetics observed with multiple degraders. Recognition of protein maturation state as a determinant of degradability may inform next□generation strategies for rational PROTAC design and improve the durability of targeted protein degradation in therapeutic settings.

## Materials and Methods

### Chemicals

Halo ligand **1** was synthesized as previously described.^25^ dBET6 and MZ-1 were obtained from MedChemExpress (Monmouth Junction, NJ). Cycloheximide (CHX) was purchased from VWR Life Science (Radnor, PA). Halo-JF549 and Halo-JF646 were obtained from Promega (Madison, WI). NucBlue was purchased from Thermo Fisher Scientific.

### Cell Lines and Culture

HEK293T cells were obtained from ATCC, and MDA-MB-468 cells were sourced from the National Cancer Institute Developmental Therapeutics Program. Both cell lines were authenticated by STR profiling and confirmed to be mycoplasma-free by PCR testing. Cells were maintained in high-glucose Dulbecco’s modified Eagle’s medium (DMEM, Thermo Fisher) supplemented with 10% FBS (Hyclone) and 10% nonessential amino acids (Thermo Fisher) at 37 °C in a 5% CO□incubator. All experiments were performed using cells within 50 passages.

### Plasmids and Antibodies

Flag-HaloTag-BRD4 was engineered as an *N*-terminal fusion comprising a Flag tag, HaloTag, and BRD4. The in-frame construct was generated by gene synthesis (Genewiz, South Plainfield, NJ) and subcloned into the third-generation lentiviral vector pLJM1 (Addgene). The sequence of HaloTag-BRD4 is provided Figure S1. The following antibodies were used: mouse anti-Flag (M2, Sigma), mouse anti-BRD4 (Cell Signaling Technology), rabbit anti-BRD4 (Fortis Life Sciences), mouse anti-GAPDH (Santa Cruz Biotechnology), and rabbit anti-Hsp90 (Cell Signaling Technology). Horseradish peroxidase (HRP)-conjugated secondary anti-mouse and anti-rabbit antibodies were obtained from Jackson Immunoresearch Laboratories.

### Lentivirus Preparation and Transduction

Lentivirus production was performed as previously described.^37, 38^ Briefly, HEK293T cells were co-transfected with the Flag-HaloTag-BRD4 lentiviral expression plasmid and packaging vectors using the calcium phosphate method (TaKaRa) to generate Flag-HaloTag-BRD4-expressing lentiviruses. MDA-MB-468 cells were then transduced with the resulting virus. The cells were selected using puromycin (0.2 μg/mL) and maintained under puromycin. During experiments, the cells were plated in puromycin-free media.

### Confocal Microscopy

This procedure is similar to the previously published method ^25^ with slight modifications. For imaging fixed cells for existing and newly synthesized HaloTag-BRD4, MDA-MB-468 cells stably expressing Flag-HaloTag-BRD4 were cultured in DMEM with 10% FBS and puromycin. Once the cells reached ⁓80% confluence, they were harvested and transferred to coverslips, which were coated with poly-D-lysine (R&D systems) for 24 h. Cells were then treated with DMSO or CHX (100 μg/mL) for one hour, followed by the addition of 10 μM Halo ligand **1** or 200 nM Halo-JF646 for one hour. The labeling media were removed, and the cells were washed six times with drug-free fresh media. The cells were incubated with fresh media containing 100 μg/mL CHX or DMSO for indicated time period. Then, the newly synthesized HaloTag-BRD4 was labelled with 200 nM Halo-JF549 for 15 minutes. The cells were fixed with 37 °C-prewarmed fixation solution at room temperature for 10 minutes. The fixation solution was 4% paraformaldehyde and 0.2 M sucrose in 1xPBS (pH 7.5). The cells were permeabilized with 0.5% triton X-100 at room temperature for 15 minutes. Then the cells were blocked in 3% BSA in PBS for 1 h at room temperature followed by with 300 nM DAPI for 10 min at room temperature. The coverslips were mounted onto glass slides. Images were taken on an inverted Zeiss LSM 980 confocal microscope.

For immunofluorescence, MDA-MB-468 cells stably expressing Flag-HaloTag-BRD4 or parental MDA-MB-468 cells were cultured in DMEM with 10% FBS. Once the cells reached ⁓80% confluence, they were harvested and transferred to coverslips, which were coated with poly-D-lysine (R&D systems) for 24 h. The cells were fixed with 4% paraformaldehyde and permeabilization for 15 minutes, the cells were blocked in 3% BSA in PBS for 1 h at room temperature followed by overnight incubation with rabbit or mouse anti-BRD4 antibody (1:1000 in PBS with 3% BSA) and mouse M2 (1:1000 in PBS with 3% BSA) at 4 °C. The next day, the cells were stained with a secondary Alexa fluor 555 conjugated anti-rabbit antibody (Invitrogen) and a secondary Alexa fluor 488 conjugated anti-mouse antibody (Jackson ImmunoResearch Laboratories) for 1 h at room temperature. The cells were further incubated with 300 nM DAPI for 10 min at room temperature. The coverslips were mounted onto glass slides. Images were taken on an inverted Zeiss LSM 980 confocal microscope.

### Live-cell dual-color monitoring of HaloTag-BRD4 degradation

MDA-MB-468 cells stably expressing HaloTag-BRD4 were plated in a 96-well glass-bottom plate (Cellvis) pretreated with poly-D-lysine (R&D systems) (2 x 10^4^ cells/well) and allowed to attach to the bottom of the plate overnight. To label existing HaloTag-BRD4, the cells were treated with 200 nM Halo-JF549 (Promega) for 1 hour, followed by washing with drug-free media 6 times to remove excess ligand. Then the cells were incubated with fresh media for 1 h to allow new protein synthesis. Subsequently, Halo-JF646 (200 nM, Promega) was added to the cells for 15 minutes to label newly synthesized HaloTag-BRD4. This was followed by a second wash step using drug-free media (6 x) to remove unbound dye. After the dual-labeling, cells were treated with DMSO or different concentrations of dBET6 or MZ-1 together with NucBlue (Invitrogen, cat # R37605). To prevent any additional labeling by residual fluorescent Halo ligand, ligand **1** (10 μM) was added concurrently with dBET6 or MZ-1. Live-cell imaging was started as soon as dBET6 or MZ-1 was added. Images were acquired every 10 minutes for 3 hours using a ZEISS Celldiscoverer 7 system, collecting Halo-JF549 (red), Halo-JF646 (far-red), and NucBlue channels. Nuclear regions were tracked and segmented based on the NucBlue channel using ZEISS arivis software, and nuclear mean intensities for the red and far-red signals were quantified. The fluorescence intensities were normalized to the signal at time zero, which was defined as 1.0. Then, the ratio of relative fluorescent intensity was calculated by the normalized-newly synthesized HaloTag-BRD4 over normalized existing HaloTag-BRD4.

### Endogenous BRD4 degradation by dBET6

Sterilized glass coverslips (Fisher Scientific) were placed in 24-well plates (Thermo Scientific) and coated with poly-D-lysine (R&D Systems). MDA-MB-468 cells were plated onto the coverslips at a density of 2×10^5^ cells per well and allowed to attach overnight. For analysis of existing BRD4, the cells were treated with CHX (100 μg/mL) for 4 hours to block protein synthesis, followed by six washes with drug-free media. Then DMSO or dBET6 at different concentrations, along with CHX, were added for another 4 h. For analysis of newly synthesized BRD4, the cells were first treated with dBET6 (100 nM) for 4 h to acutely deplete existing BRD4. Following this treatment, the cells were washed six time with drug-free media to remove residual dBET6. Then, the cells were incubated with DMSO or dBET6 at different concentrations for another 4 h. At the end of treatment, the cells were fixed with 4% paraformaldehyde and permeabilized with 0.1% Triton X-100. Following fixation and permeabilization, the cells were blocked in 3% BSA in PBS for one hour at room temperature. The blocked cells were then incubated with mouse anti-BRD4 primary antibody overnight. The following day, the cells were incubated with Alexa Fluor 555-conjugated anti-mouse secondary antibody. The cells were counterstained with 300 nM DAPI for 10 minutes at room temperature. Fixed cell immunofluorescence imaging was performed using Zeiss LSM980 confocal microcopy.

An image analysis pipeline was created in the ZEN Image Analysis module. First, the DAPI channel was assigned as the segmentation channel. Then global thresholding was applied to the DAPI signal to identify nuclear boundaries, and threshold values were interactively optimized using the histogram-based thresholding tool. Watershed separation was applied to resolve adjacent nuclei, and hole filling was enabled. A region filter was then applied to exclude objects outside the expected nuclear size range. For each segmented nucleus, ZEN automatically generated nuclear masks that were applied to all imaging channels. Using the Features panel, the “Intensity: Mean Value” feature was selected. During batch analysis, ZEN computed per-nucleus mean fluorescence intensity for the red channels based on the DAPI-defined nuclear masks. These measurements were exported as analysis tables and used for downstream quantification. Intensities in the new BRD4 assay were normalized to the DMSO control, whereas intensities in the existing BRD4 assay were normalized to the CHX + DMSO condition.

### Western blot

MDA-MB-468 cells were plated into 6-well plates and allowed to attach to the bottom of the plates for overnight. Then the cells were treated dBET6 for 24, 48 or 72 h. Then the cells were collected by mechanical scraping. The cell pellets were washed twice with ice-cold PBS. The cell pellets were lysed in lysis buffer A (50 mM Tris, 5 mM EDTA, 150 mM NaCl, 1 mM DTT, 0.5% Nonidet P-40, pH 8.0) supplemented with protease inhibitor cocktail (Roche) and 1 mM PMSF. The concentration of the cleared lysates was determined using Dye Reagent Concentrate (Bio-Rad). Then equal amount of protein was loaded onto pre-cast 4-20% SDS-PAGE (Bio-Rad) for electrophoresis. The proteins were transferred to a nitrocellulose membrane using Trans Blot^®^ Turbo^TM^ Transfer system (Bio-Rad). The membrane was blocked in 5% non-fat dry milk for 1 h at room temperature followed by incubation with a primary antibody for overnight at 4 °C. The membrane was washed and then incubated with horseradish peroxidase (HRP)-conjugated secondary antibodies for bioluminescence imaging using ChemiDoc^MP^ (Bio-Rad).

### Statistical analysis

Pair wise comparisons were performed using student *t*-test in Microsoft Excel (Version 16). The time-course analyses were performed using IBM SPSS Statistics (version 30.0.0, IBM Corp., Armonk, NY, USA). A mixed-model repeated-measures ANOVA was conducted to evaluate signal dynamics across time. Treatments and signal type (red and far-red) were specified as between-group factors, and time (18 time points over a 3-hour imaging period) was included as a within-group factor. A *P* value of less than 0.05 was deemed significant.

## Acknowledgements

We appreciate the financial supports provided by National Institutes of Health (R01GM122820, R01CA278058, R01CA245964) and Oregon Health & Science University (OHSU) School of Medicine. We thank Professor Byung Park (OHSU) for expert help on statistical analysis and Dr. Felice Kelly (OHSU) for technical support on live cell imaging. We thank OHSU Biophysical Shared Resources Core for providing various biophysical instrument to support this work. We appreciate OHSU Integrated genomics core for authenticating the cell lines through STR profiling.

