## Supplementary for "Protein Age Bias in Target Degradation by PROTACs"

>Flag-HaloTag-BRD4

MDYKDHDGDYKDHDIDYKDDDDKASAEIGTGFPFDPHYVEVLGERMHYVDVGPRDGTPVLFLHGNPTSSYVWRNIIPHVAPTHRCIAPDLIGMGKSDKPDLGYFFDDHVRFMDAFIEALGLEEVVLVIHDWGSALGFHWAKRNPERVKGIAFMEFIRPIPTWDEWPEFARETFQAFRTTDVGRKLIIDQNVFIEGTLPMGVVRPLTEVEMDHYREPFLNPVDREPLWRFPNELPIAGEPANIVALVEEYMDWLHQSPVPKLLFWGTPGVLIPPAEAARLAKSLPNCKAVDIGPGLNLLQEDNPDLIGSEIARWLSTLEISGSRMSAESGPGTRLRNLPVMGDGLETSQMSTTQAQAQPQPANAASTNPPPPETSNPNKPKRQTNQLQYLLRVVLKTLWKHQFAWPFQQPVDAVKLNLPDYYKIIKTPMDMGTIKKRLENNYYWNAQECIQDFNTMFTNCYIYNKPGDDIVLMAEALEKLFLQKINELPTEETEIMIVQAKGRGRGRKETGTAKPGVSTVPNTTQASTPPQTQTPQPNPPPVQATPHPFPAVTPDLIVQTPVMTVVPPQPLQTPPPVPPQPQPPPAPAPQPVQSHPPIIAATPQPVKTKKGVKRKADTTTPTTIDPIHEPPSLPPEPKTTKLGQRRESSRPVKPPKKDVPDSQQHPAPEKSSKVSEQLKCCSGILKEMFAKKHAAYAWPFYKPVDVEALGLHDYCDIIKHPMDMSTIKSKLEAREYRDAQEFGADVRLMFSNCYKYNPPDHEVVAMARKLQDVFEMRFAKMPDEPEEPVVAVSSPAVPPPTKVVAPPSSSDSSSDSSSDSDSSTDDSEEERAQRLAELQEQLKAVHEQLAALSQPQQNKPKKKEKDKKEKKKEKHKRKEEVEENKKSKAKEPPPKKTKKNNSSNSNVSKKEPAPMKSKPPPTYESEEEDKCKPMSYEEKRQLSLDINKLPGEKLGRVVHIIQSREPSLKNSNPDEIEIDFETLKPSTLRELERYVTSCLRKKRKPQAEKVDVIAGSSKMKGFSSSESESSSESSSSDSEDSETEMAPKSKKKGHPGREQKKHHHHHHQQMQQAPAPVPQQPPPPPQQPPPPPPPQQQQQPPPPPPPPSMPQQAAPAMKSSPPPFIATQVPVLEPQLPGSVFDPIGHFTQPILHLPQPELPPHLPQPPEHSTPPHLNQHAVVSPPALHNALPQQPSRPSNRAAALPPKPARPPAVSPALTQTPLLPQPPMAQPPQVLLEDEEPPAPPLTSMQMQLYLQQLQKVQPPTPLLPSVKVQSQPPPPLPPPPHPSVQQQLQQQPPPPPPPQPQPPPQQQHQPPPRPVHLQPMQFSTHIQQPPPPQGQQPPHPPPGQQPPPPQPAKPQQVIQHHHSPRHHKSDPYSTGHLREAPSPLMIHSPQMSQFQSLTHQSPPQQNVQPKKQELRAASVVQPQPLVVVKEEKIHSPIIRSEPFSPSLRPEPPKHPESIKAPVHLPQRPEMKPVDVGRPVIRPPEQNAPPPGAPDKDKQKQEPKTPVAPKKDLKIKNMGSWASLVQKHPTTPSSTAKSSSDSFEQFRRAAREKEEREKALKAQAEHAEKEKERLRQERMRSREDEDALEQARRAHEEARRRQEQQQQQRQEQQQQQQQQAAAVAAAATPQAQSSQPQSMLDQQRELARKREQERRRREAMAATIDMNFQSDLLSIFEENLF

**Figure S1**. The protein sequence of Flag-HaloTag-BRD4 used in this study.

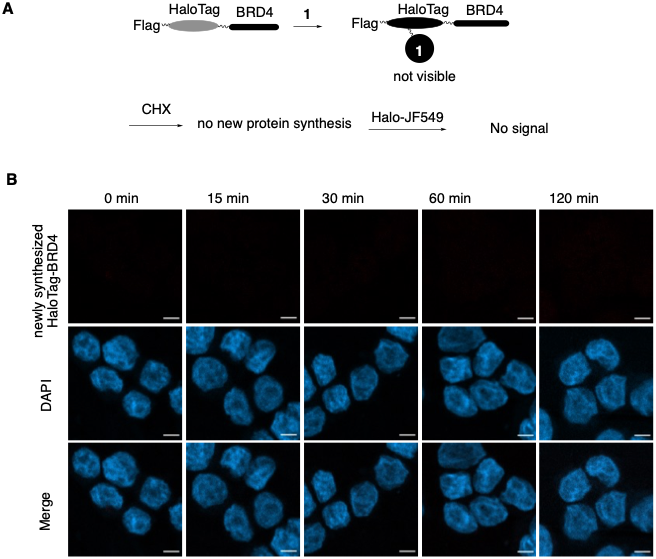

**Figure S2.** CHX blocks new synthesis of HaloTag-BRD4. (A) Experimental design to investigate CHX’s effect on new HaloTag-BRD4 synthesis. (B) Representative images of lack of newly synthesized HaloTag-BRD4 to be labeled by Halo-JF549. MDA-MB-468 cells with HaloTag-BRD4 were treated with Halo ligand **1** for 30 min. Then the cells were treated with CHX for different periods of time, when Halo-JF549 was added for another 10 min. The nuclei were counter stained with DAPI. Scale bars are 5 μm.

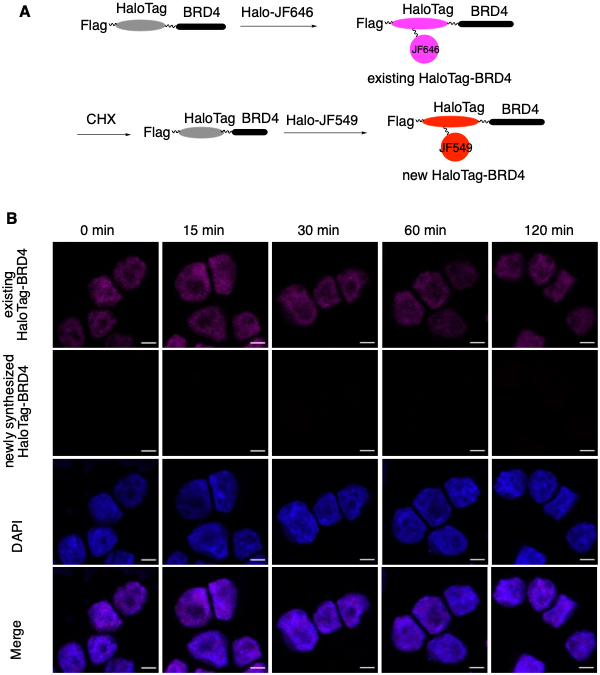

**Figure S3.** _CG_-SLENP for dual labeling of existing and newly synthesized HaloTag-BRD4. (A) A schematic diagram to illustrate the dual-labeling _CG_-SLENP strategy to label existing and newly synthesized HaloTag-BRD4 in the same cells. (B) Representative confocal micrographs of existing and newly synthesized HaloTag-BRD4. MDA-MB-468 cells expressing HaloTag-BRD4 were treated with Halo-JF646 for 1 h, when the ligand was removed. The cells were then treated with CHX for indicated time period, when the cells were treated with further Halo-JF549 for 10 min. The cells were then fixed and permeabilized. The cell nuclei were counterstained with DAPI. Scale bars are 5 μm.

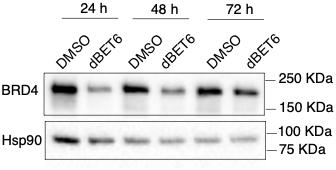

**Figure S4.** dBET6 induced degradation of BRD4. MDA-MB-468 cells were treated with DMSO or dBET6 for indicated time periods. The cells were then collected, and the resulting cell lysates were analyzed by western blot using indicated antibodies.

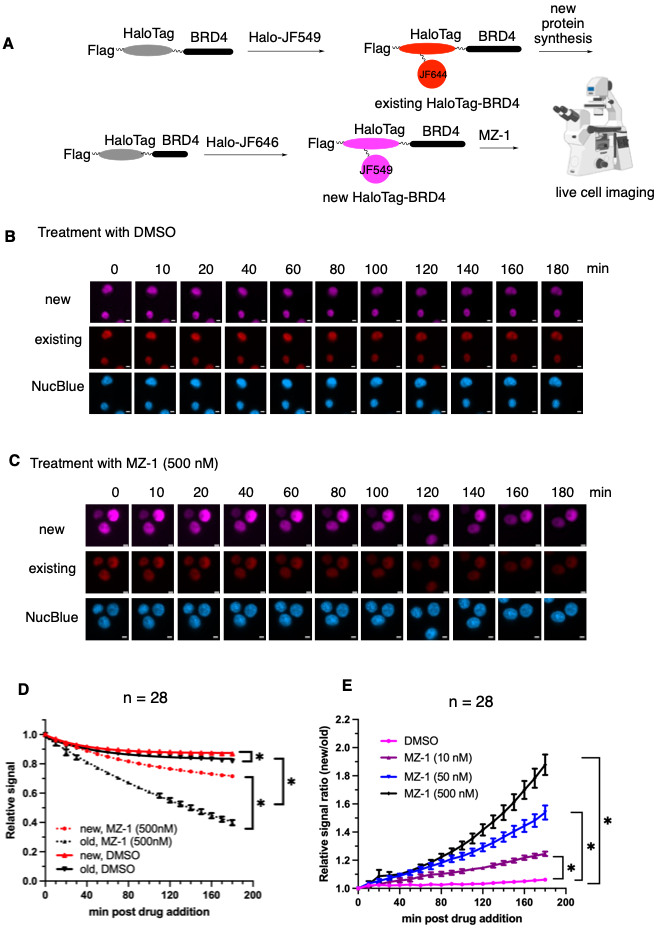

**Figure S5. MZ-1 preferentially degrades existing HaloTag-BRD4 over newly synthesized HaloTag-BRD4.** (A) A schematic diagram to illustrate dual-labeling by _CG_-SLENP to investigate the degradation effects of MZ-1 on existing and newly synthesized HaloTag-BRD4 in live cells. (B, C) Representative live cell micrographs of cells treated with DMSO (C) or MZ-1 (500 nM, C). Existing HaloTag-BRD4 was labeled with Halo-JF549 while the newly synthesized HaloTag-BRD4 was labeled with Halo-JF646 (far-red). Upon labeling, the cells were then treated with DMSO or different concentrations of MZ-1 for live-cell imaging. The images were acquired every 10 minutes for 3 hours in the red, far-red, and NucBlue channels. (D) Time-dependent degradation of HaloTag-BRD4 by MZ-1. The existing and newly synthesized HaloTag-BRD4 signals from images in (B) and (C) were quantified. Data are presented as mean ± SEM (n = 28). **P*<0.01. (E) MZ-1 dose-dependently induced preferential degradation of existing HaloTag-BRD4. The data in (D) were replotted by dividing newly synthesized HaloTag-BRD4 to the existing HaloTag-BRD4. These divided values were normalized to 1.0 at time zero. Data are presented as mean ± SEM (n = 28). **P*<0.01.
